# Gobies (Perciformes: Gobiidae) in Bolinao, northwestern Philippines

**DOI:** 10.1101/2020.03.20.999722

**Authors:** Klaus M. Stiefel, Dana P. Manogan, Patrick C. Cabaitan

**Affiliations:** The Marine Science Institute, University of the Philippines, Diliman, Quezon City, Philippines; Neurolinx Research Institute, La Jolla, CA, USA

**Keywords:** Bolinao, Coral Reef, Fish, Gobiidae, Philippines

## Abstract

We conducted a visual and photographic survey of the gobiidae in the Bolinao area of the Philipines, located on the western tip of the Lingayen gulf, on the west coast of Luzon island. We identified a total of 40 species, of which 18 are shrimp-associated. One species found (*Myersina lachneri*) constitutes a range expansion into the Philippines. This number of species is in the expected range compared to other studies of marine goby faunae in the coral triangle, despite the significant anthropogenic pressures onto the marine ecosystem in the surveyed area.

## INTRODUCTION

Gobies (Perciformes: Gobiidae) are the largest family of marine fishes, with over 1800 known species (1834 listed on Fishbase, Froese & Pauly, 2010). In tropical coastal ecosystems, gobies constitute a significant fraction of all fish species. Many species of gobies are cryptic, living as epibionts on corals and sponges, or highly camouflaged in the sand. About 120 species of marine gobies also live in a symbiotic relationship with alpheid shrimp, with which they share a burrow excavated by the shrimp. In mangrove areas, mudskippers of the genus *Pteriophtalmus* are amphibious and venture out onto the mud between the mangrove roots. Most species of gobies are small, with *Schindleria brevipinis* possibly the smallest known vertebrate (7 mm adult length, Watson & Walker, 2004). Generally, knowledge of the gobiid fauna provides a valuable window into the fish diversity of a location.

## MATERIALS & METHODS

We used visual and photographic identification by two or three observers during SCUBA dives to survey an area of about 75 km^2^ around Santiago island east of Bolinao on the western edge of the Lingayen gulf (Fig. 1; 16° 24’ 32” North, 119° 56 ‘13” East), northwestern Philippines.

**Fig. 1.**
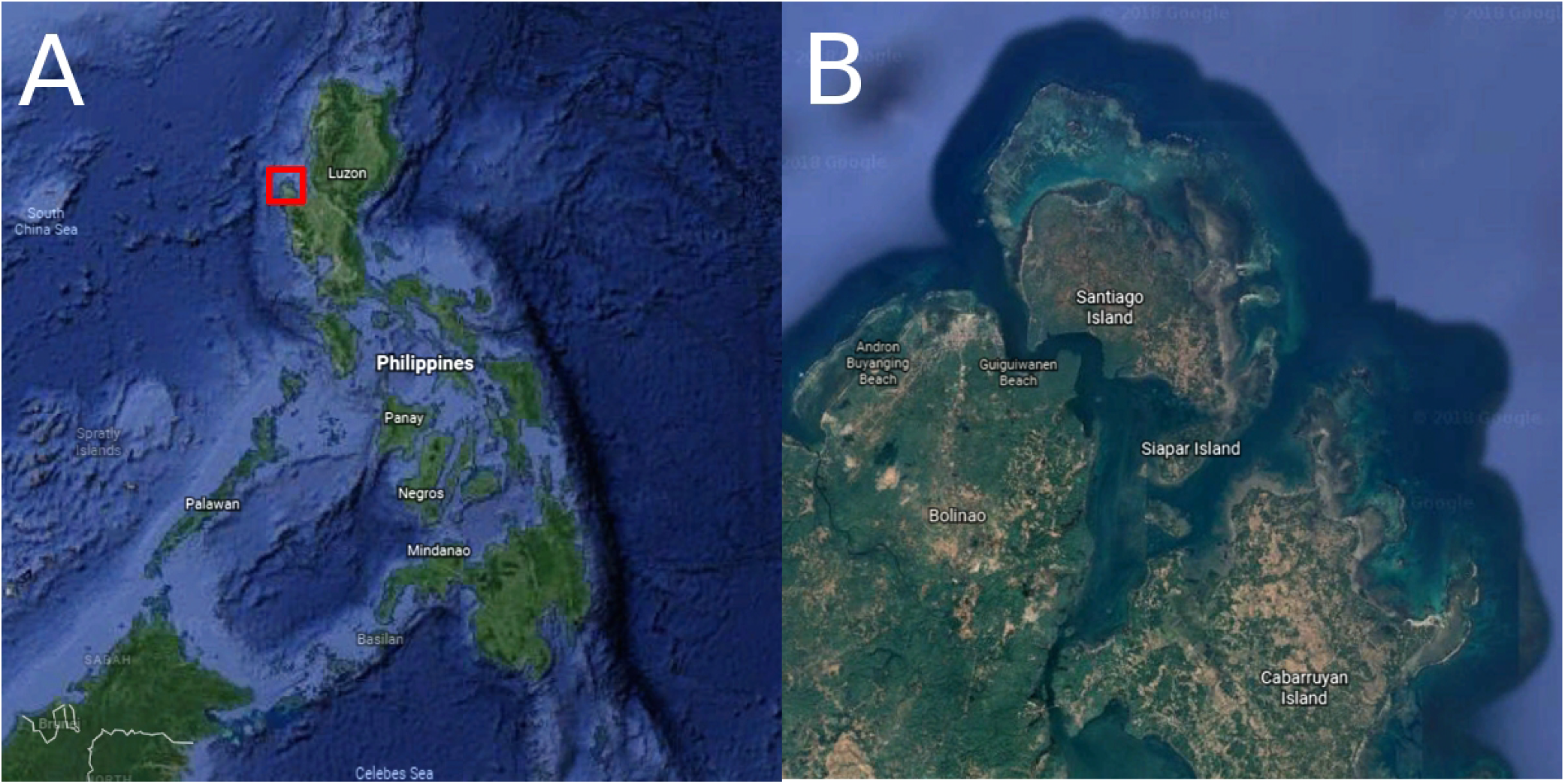
Survey area around Santiago island (B) in the Bolinao region, Pangasinan province, Luzon island, Philippines (A).

The sampling sites were between 1 and 35 meters deep, with the majority of species found shallower than 18 meters. Habitats included coral reefs, sandy areas adjacent to reefs, sandy/silty areas not in proximity to coral reefs, as well as seagrass areas. An unusual sampling site is the giant clam ocean hatchery of the University of the Philippines Marine Science Institute, which is home to several thousand giant clams (*Tridacna* spp.). These clams provide 3-dimensional structure similar to a coral reef. We excluded esturine, brackish water and freshwater habitats.

The anthropogenic disturbances of the marine environment in the western Lingayen gulf are significant, with fish farms introducing a significant amount of nutrients, and strong artisanal and large-scale fishing operations depleting fish stocks (McManus, 1992; Campos et al., 1994). The drop in water quality caused by the fish farming on the west side of Santiago Island has led to coral reef degradation and in some spots to a conversion of former reefs to silty areas devoid of corals (Cabaitan et al., 2016).

## RESULTS

We found a total of 40 species of gobiidae, of which 18 were shrimp-associated, 7 were coral epibionts at least part of the time, 14 were sand-living, 1 found in rubble and 1 in rock crevices (Table 1, Fig. 2).

**Table 1.**
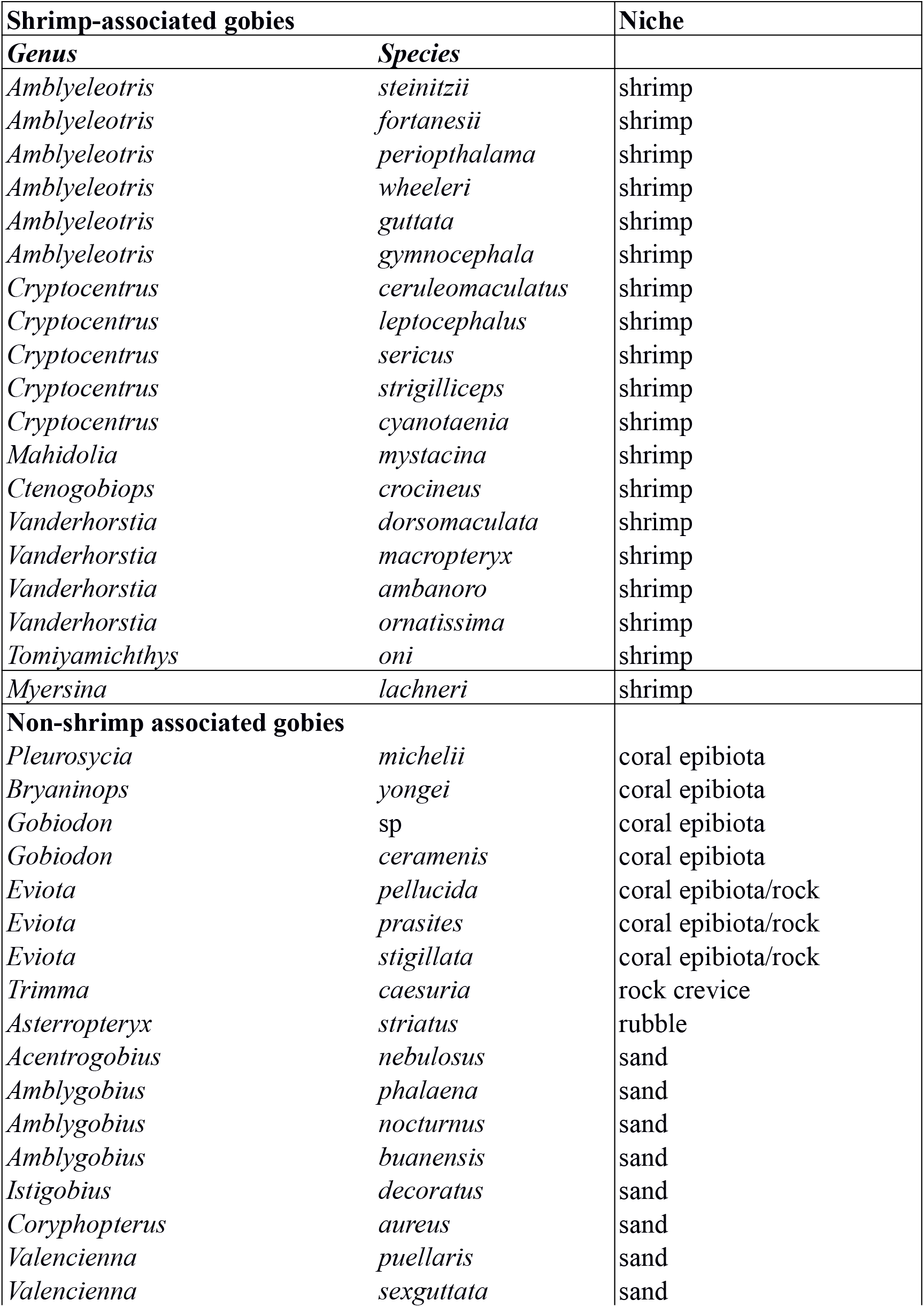
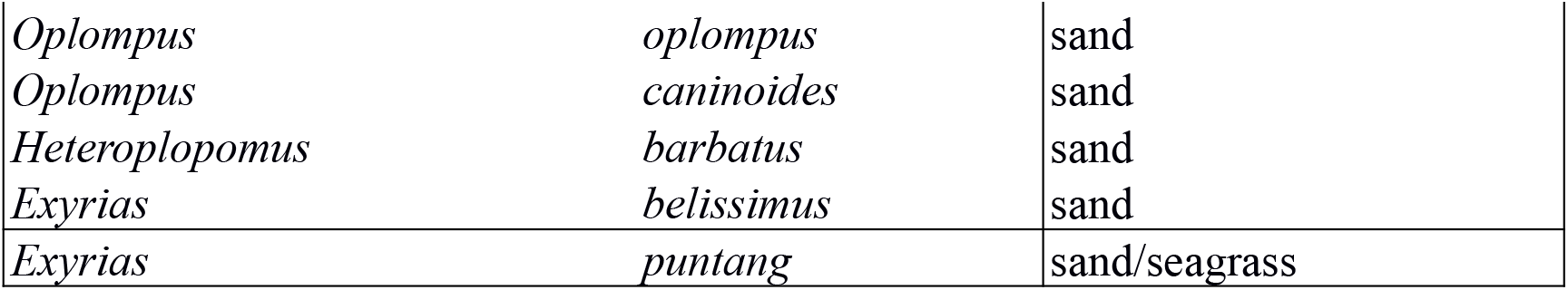
All gobiid species recorded in the vicinity of Bolinao, northwestern Philippines.

**Fig. 2.**
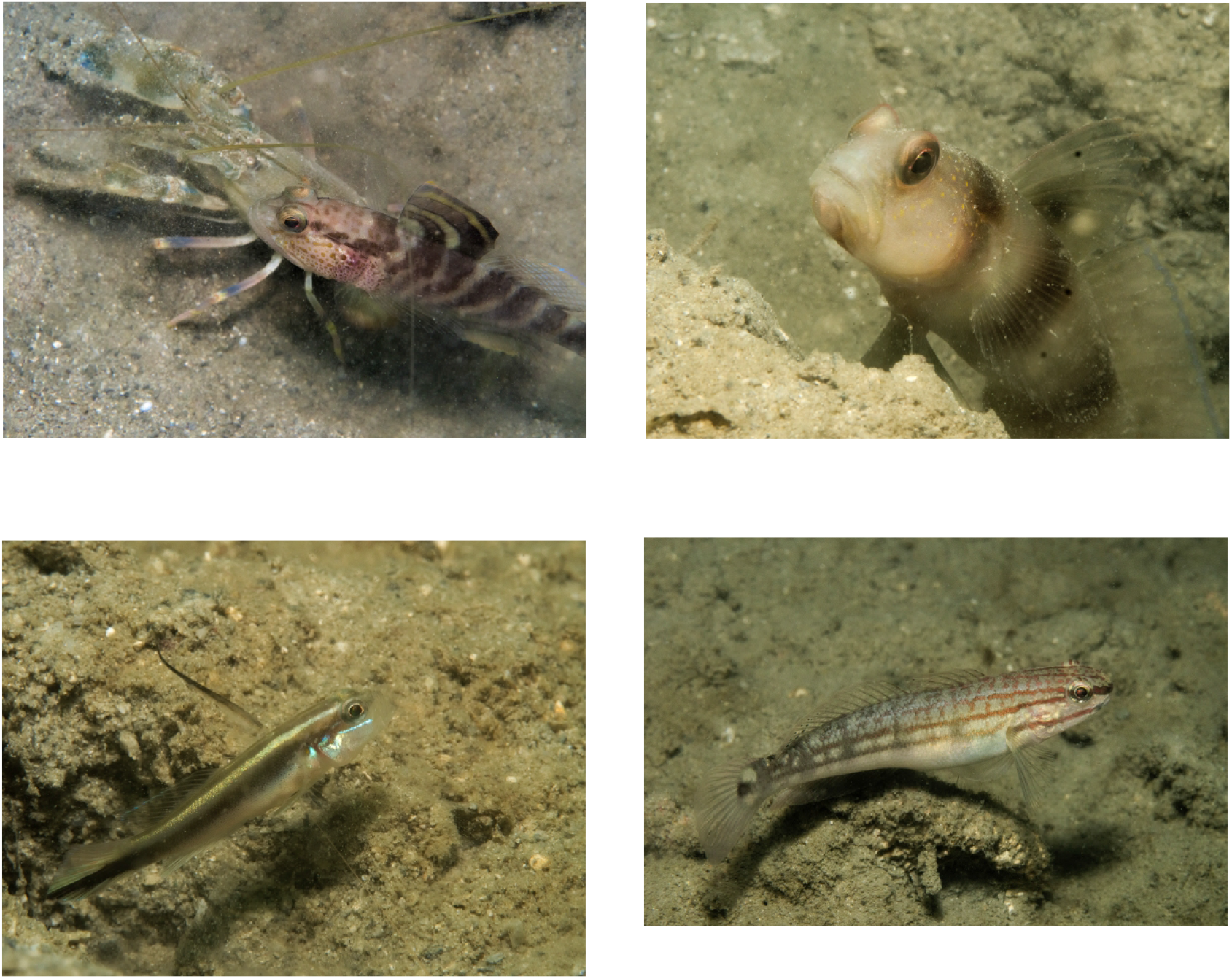
Photographs of several species encountered in the Bolinao area (top to bottom, left to right): *Mahidolia mystacina, Amblyeleotris fortanesii, Myersina lachneri, Amblygobius buanensis.*

We found individuals of all but two species (*Gobiodon* sp.*, Gobiodon ceramenis*) multiple times, indicating that while our sampling of the goby species in the area might not be complete, we had found all but the most rare species.

*Gobiodon* sp., a yellow fish with orange facial markings, is likely the undescribed species listed in Allen & Erdmann (2012). *Myersina lachneri* (Hoese & Lubbock, 1982) is a range expansion for the Philippines, having previously only been reported from Papua New Guinea and Indonesia (Allen & Adrim, 2003).

Photographs of 31 of the 41 described species are available here: https://www.flickr.com/photos/pacificklaus/sets/72157685611197132

Video footage of several species featured in the photographs, and two more are available here: https://www.youtube.com/watch?v=Q4KMPjV0qSg

## DISCUSSION

Our survey found 40 species of gobies in an area of about 75 km^2^. This is close to an expected number of species compared to other surveys of gobbiidae in the Indo-Pacific (see Fig. 3 for a species-area plot as the basis for this prediction). Surveying an area of a comparable size (40 km^2^), Depczynski & Bellwood (2005) found 30 species around Lizard Island in the GBR.

**Fig. 3.**
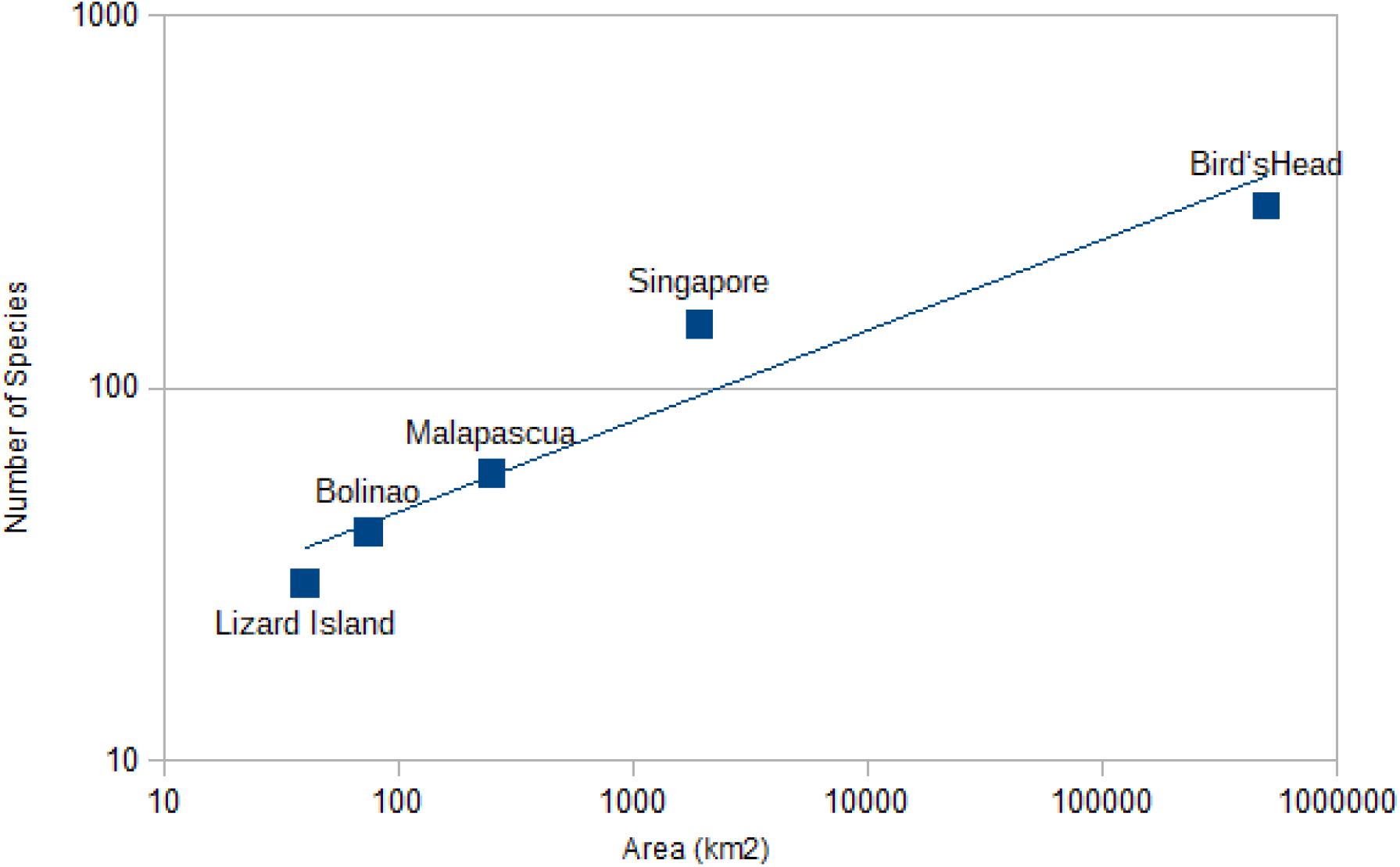
Species – area relationship for marine gobies. Plotted are the number of species against the estimated survey area from this study (~ 75 km^2^, 40 species), a study of the gobies of Lizard Island (~ 40 km^2^, 30 species), of Malapascua, Cebu province, Philippines (~ 250 km^2^, 59 species), of Singapore (~ 1925 km^2^, 149 species) and the Papuan Bird’s Head Peninsula (~50 000 km^2^, 308 species).

The goby fauna in the western Lingayen near Bolinao gulf is likely determined by the physical conditions as well as by anthropogenic disturbance. The area lacks deep walls which are habitats for hovering gobies (such as *Trimma tevegae*), which are hence absent from the area. Additionally, the eastern side of Santiago island is swept by powerful currents, known to limit the occurrence of small marine fishes (Depczynski & Bellwood, 2005). The low number of gobiid epibionts is most likely a consequence of the limited coral cover and diversity. This might partially be a consequence of the severe anthropogenic stresses to marine habitats in the region.

Nevertheless, a gobiid fauna of 40 species, close to the expected value, indicates that small, often cryptic, fishes low in the food web could be less likely affected by anthropogenic disturbances than larger species.

## ACKNOWLEDGMENTS

We would like to thank our colleagues at the University of the Philippines, Diliman, Marine Science Institute, especially Renato Adolfo for help in sampling, Dr. Cecilia Conaco and Timothy Quimpo for helpful discussion. We also thank Andreas Völkers and Dr. Brett Tibbatts for help with fish identification, and Dr. Rene Abesamis for discussion and pointers to the literature.

